# The Effects of vibrotactile stimulation of the upper extremity on sensation and perception

**DOI:** 10.1101/2024.05.02.592163

**Authors:** Abeer Abdel Khaleq, Yash More, Brody Skaufel, Mazen Al Borno

## Abstract

Vibrotactile stimulation has applications in a variety of fields, including medicine, virtual reality, and human-computer interaction. Eccentric Rotating Mass (ERM) vibrating motors are widely used in wearable haptic devices for their small size, low cost, and low energy features. The effect of ERM motor vibrations on upper extremity sensation and perception have not been thoroughly studied previously, which is important to design better wearable haptic devices. We conducted experiments with vibrotactile stimulation on 15 healthy participants. Eight motors were placed on a consistent set of muscles on the upper extremity and one motor was placed on the index finger. We found a significant correlation between voltage and sensation intensity (r=0.39). However, we did not find a significant aggregate-level correlation on the perceived pleasantness of the simulation. The sensation intensity varied based on the location of the vibration on the upper extremity (with the lowest intensity on the triceps brachii and brachialis) and slightly decreased (5.9 ± 2.9 %) when participants performed reaching movements. When a single motor was vibrating, the participants’ accuracy at identifying the motor without visual feedback increased as voltage increased, reaching up to 81.4 ± 14.2 %. When we stimulated three muscles simultaneously, we found that most participants were able to identify only two out of three vibrating motors (41.7 ± 32.3 %). Our findings can help tune stimulation parameters in human-machine interaction applications.

## Introduction

Vibrotactile feedback is a mechanical stimulation that is produced with actuators placed on the skin [1]. This form of stimulation has applications in gaming and virtual reality [2], movement training [3], rehabilitation after stroke [4] and neuromodulation in Parkinson’s disease [5]. For haptic feedback with vibrotactile stimulation to advance in these areas, it becomes necessary to understand how different parameters impact how people interact with the stimulation. Stimulating the upper extremity is interesting for applications like upper extremity stroke rehabilitation (i.e., hemiparesis) and teaching new motor skills like playing a musical instrument.

How different vibrotactile stimulation parameters applied on the upper extremity impact sensation and perception have not been previously investigated. In this work, we present a study on 15 healthy participants to analyze the effects of changing the vibration signal voltage on sensation and perception. Our contributions include 1) an analysis of sensation intensity and perceived pleasantness when changing the voltage stimulation parameter across the upper extremity; 2) a comparison of sensation intensity when the upper extremity is still and when it is actively in movement; 3) an analysis of how accurately participants can identify the vibrating motors or the stimulated muscles from tactile feedback.

## Related work

Vibrotactile feedback has shown to be effective for delivering tactile cues as their small size enable them to be embedded in light weight garments that do not hinder the movement of the participant [3]. This could help in developing tactile motion guidance for motor learning or rehabilitation therapy, so that participants can practice motions on their own without the presence of a coach or therapist. Previous work has studied the effects of some vibrotactile stimulation parameters in older adults with and without a history of stroke [1]. However, stimulation was limited to the hand and the forearm. Gtat et al. [10] found that a pulse width of 15ms had an average perceptibility of 50%, thus making it the absolute detection threshold for the average participant. Ng et al. [7] found that high amplitude and low frequency stimuli were perceived more intensely than high frequency and low amplitude stimuli. Our study focuses on the effects of changing the voltage parameter of ERM motors when the stimulation is applied on the shoulder, upper arm, lower arm, and index finger. Other studies have focused on understanding the effect of the stimulation pattern on controlling an upper limb prosthetic [6]. They have found that the stimulation pattern had no significant effect on sensation intensity. Furthermore, they did not observe a significant difference between the effects of amplitude and frequency on sensation intensity. In our work, we use ERM motors where frequency and amplitude are linked. We are interested in studying the effects of changing the voltage on sensation, perception, and motor identification in healthy adults.

It is known that tactile sensation in the hand decreases as it is in movement [8] and our work examines whether this effect is also observed throughout the upper extremity. Bark et al. [3] evaluated the effect of vibrotactile feedback on the learning of arm motions. The vibrating motors were placed on the forearm without regard to placing the motors consistently on the same muscles. In our work, we place the motors on important muscles in the upper extremity, which can help to study the effects of vibrotactile stimulation on teaching new muscle coordination patterns. Shah et al. [9] developed a study on distinguishing stimuli thresholds in the upper extremity using both sequential and simultaneous vibration frequencies. They observed that that discrimination threshold in the front of the forearm was on average 10 Hz lower than the threshold for the back of the forearm. Their results showed that sequentially delivered stimuli were identified more accurately than simultaneously delivered stimuli. We investigate in our study if participants can identify simultaneous vibrating motors in different regions of the upper extremity and at what threshold values.

## Vibrotactile Stimulation System

We present in this section the vibrotactile stimulation system developed for this study. The system consists of a microcontroller from Arduino (Mega 2560) that has 15 digital pins which can be used for the pulse width modulation (PWM) signal. The maximum voltage value of the digital pins is 5V. The pins are connected to ERM vibrating motors from Seeed Technology (Model # 316040001). We use ERM vibrating motors where vibration voltage and frequency are linked [11].

We measured the frequencies of the motors using an accelerometer and the Fourier transform. In Fig. 1, we show how the measured frequencies and voltages vary with our ERM motors. We place 8 motors on specific muscles on the shoulder, upper arm, lower arm and 1 vibrating motor on the index finger. Fig. 2 provides a general description of the system components and shows the placement of the motors on the upper extremity muscles and on the index finger. The control unit of the vibrotactile system is developed in MATLAB. The vibrotactile signal is generated with PWM sent to the Arduino digital pins connected to the vibrating motors. We control the vibrating signal voltage and duration in real-time. We record the participant’s verbal response to the changes in these parameters. The control unit communicates with the motors individually via serial communication. We change the voltage value entered to the motors by changing the PWM duty cycle using the writePWMDutyCycle function in MATLAB [12].

**Fig. 1.**
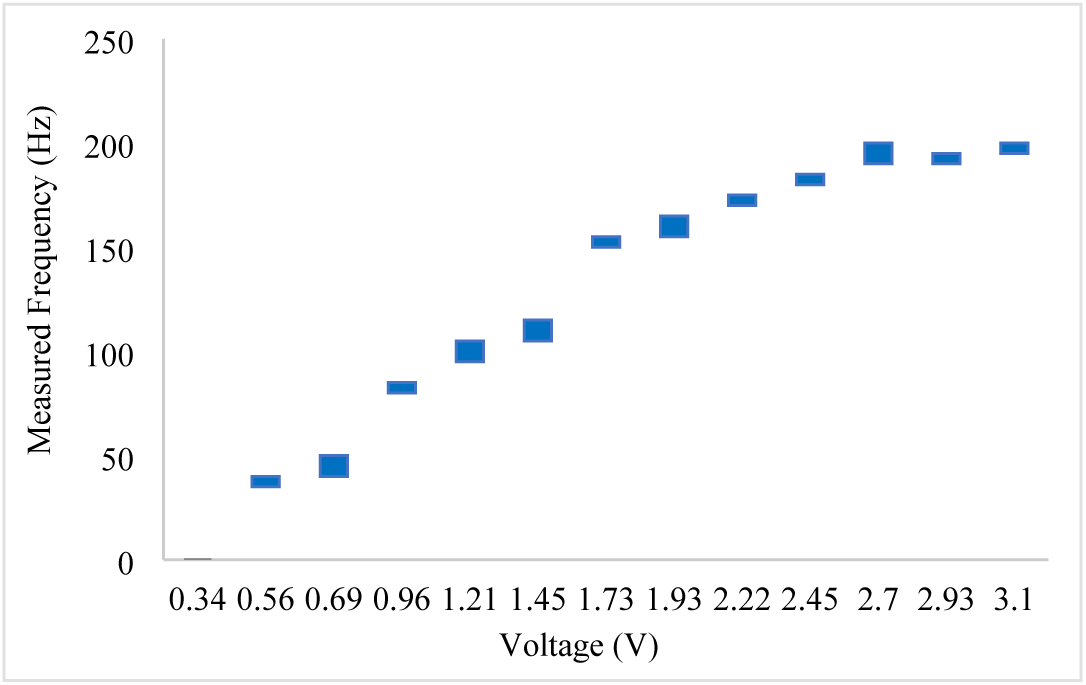
The relationship between measured voltage and frequency with the ERM motors.

**Fig. 2.**
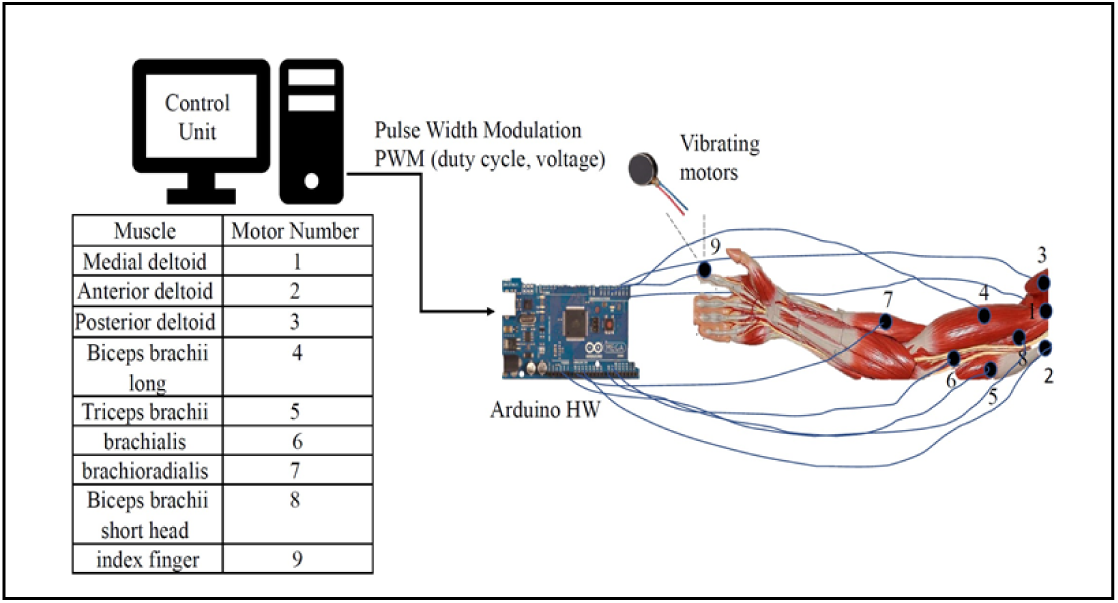
The vibrotactile stimulation system.

We observe how both frequency and voltage are linked: as we increase voltage, frequency increases. The bars denote the range of frequencies measured.

The system consists of two main components: the Arduino Mega device with attached vibrating motors (numbered from 1-9) and the control unit software that sends the voltage values and records the subject feedback in real-time.

## Sensation and perception studies and protocol

We conducted three separate studies on each participant to investigate how they sense and perceive the stimulation when the upper extremity is still, how they sense the stimulation during movement, and to characterize their conscious awareness of which motor is vibrating. Throughout the paper we will use sensation to refer to the level of sensation intensity felt on a scale from 1 to 4 (1. not felt, 2. low, 3. medium, 4. high). We will use the term perception to refer to the perceived pleasantness of the stimulation. This will be rated on a scale from 1 to 3 (1. unpleasant, 2. neutral, 3. pleasant). One of our objectives is to determine how changing the stimuli’s voltage affects sensation intensity and perceived pleasantness. The studies were conducted on 15 healthy adults (13 males and 2 females) with an average age of 26.6 ± 5.2 years. Each participant participated in the three studies on the same day during consecutive sessions. Participants were given an opportunity to take a break between sessions. The total time for the experiments was about an hour per participant. The sample size was chosen based on similar behavioral studies [1]. Participants understood and consented to the protocol approved by the Institutional Review Board of the University.

### Initial setup

At the start of the session, we asked the participant to be seated and we placed the vibrotactile device on their dominant upper extremity. The vibrating motors were placed directly on the skin, and they were attached by an elastic tape. The motors were placed on the belly of the following muscles for all participants: anterior deltoid, medial deltoid, posterior deltoid, triceps brachii, brachioradialis, brachialis, biceps brachii long head, and biceps brachii short head. An additional motor was placed on the index finger. We start the session by applying random vibrotactile stimulations for two minutes with different voltage values to get the participants accustomed to the vibrations.

## Still upper extremity study

The objective of this study is to analyze how sensation and perception are affected by changes in vibrotactile voltage. We are also investigating whether participants can identify the locations on the upper extremity of the vibrating motors, without visual feedback.

### Protocol

One motor (out of nine) was chosen at random and stimulated for 30 cycles, which lasted for about 10 s. We generated various voltage values as shown in Table 1. We applied voltage values chosen randomly from Table 1 on each motor separately. After each motor vibrates, the participants were asked to rate the sensation intensity and the perceived pleasantness on a scale from 1 to 4 and 1 to 3, respectively.

**Table 1.** Values of generated voltage measured on the motors.

| Duty Cycle Input out of 255 (DC) | Measured Voltage Output (V) |
| --- | --- |
| 51 | 0.55 – 0.56 |
| 64 | 0.68 – 0.69 |
| 128 | 1.26 – 1.30 |
| 255 | 2.54 – 2.57 |

Participants were also asked if they could identify the location of the vibrating motor precisely, vaguely, or not at all, and without looking at the upper extremity device. If the response was vaguely, they were asked to identify the upper extremity region (i.e., the shoulder, upper arm, or lower arm).

## Moving upper extremity study

In this study, we aim to determine the effects of movement on sensation and motor identification across the upper extremity.

### Protocol

Participants performed reaching movements back-and-forth at their chosen pace for the duration of the study, which was less than 7 minutes. Different voltage values chosen randomly from Table 1 were applied on the medial deltoid, triceps brachii, biceps short head and brachioradialis. A subset of the motors was chosen compared to the still study to reduce the duration of the experiment and to ensure participants do not get tired.

## Multiple vibrating motors study

In this last study, we vibrated multiple motors simultaneously across the upper extremity with the goal of determining whether participants can identify which motors are vibrating, or whether we observe differences between certain upper extremity muscles and regions.

### Protocol

In this study, three motors vibrated simultaneously for 60 seconds, and participants were asked to identify which motors are vibrating. The three motors were chosen from one of the following sets: 1) anterior deltoid, biceps brachii long head, triceps brachii; 2) the medial deltoid, posterior deltoid, biceps brachii short head; 3) brachioradialis, brachialis, biceps brachii long; and 4) triceps brachii, brachioradialis, biceps brachii short head. The sets were chosen so that the muscles span different regions in the upper extremity. For this experiment, we applied a voltage of 5 V, which corresponds to an average of 2.56 V ± 0.02 measured voltage, on all motors. After each set of motors stops vibrating, the participant was asked to evaluate the motor locations (i.e., precisely, vaguely, or not at all). If they responded vaguely, then they were asked to identify the regions where the motors were vibrating (i.e., shoulder, upper arm, or lower arm). During this experiment, participants were asked not to look at their upper extremity to ensure they do not identify the vibrating motors with visual feedback instead of tactile feedback.

## Results and analysis

In the sub-sections below, we present our results along with an analysis for the three study outcomes.

### Sensation and perception analysis in the still upper extremity

Our first study focused on the effect of changing voltage on sensation and perceived pleasantness when the participant’s upper extremity is at rest. The total dataset has 1224 records. We grouped the results based on the average measured voltage value into LOW (0.63 V ± 0.01), MEDUIM (1.28 V ± 0.03), and HIGH (2.56 V ± 0.02). The motor vibration frequencies associated with these voltage values may stimulate a range of sensory receptors (especially as the motors ramp up and down), including the Pacinian corpuscles cutaneous mechanoreceptors preferentially at 200 Hz [7] and the muscle spindle receptors at 80 Hz [18]. In this work, we consider p values less than 0.05 to be statistically significant. We found a moderate to strong positive correlation between voltage and sensation intensity (Pearson correlation coefficient of r=0.39 with p<0.05), We found no significant correlation between perceived pleasantness and voltage. We can observe this in Fig. 3, where the perception level remains stable in the neutral range with an average value of 2.2 ± 0.05 and does not incur significant changes when increasing voltage. Sensation intensity increases from 1.46 ± 0.54 (at low voltage) to 3.37 ± 0.87 (at high voltage) on average. The average increase in sensation intensity is 53.3 ± 2.9 %.

**Fig. 3.**
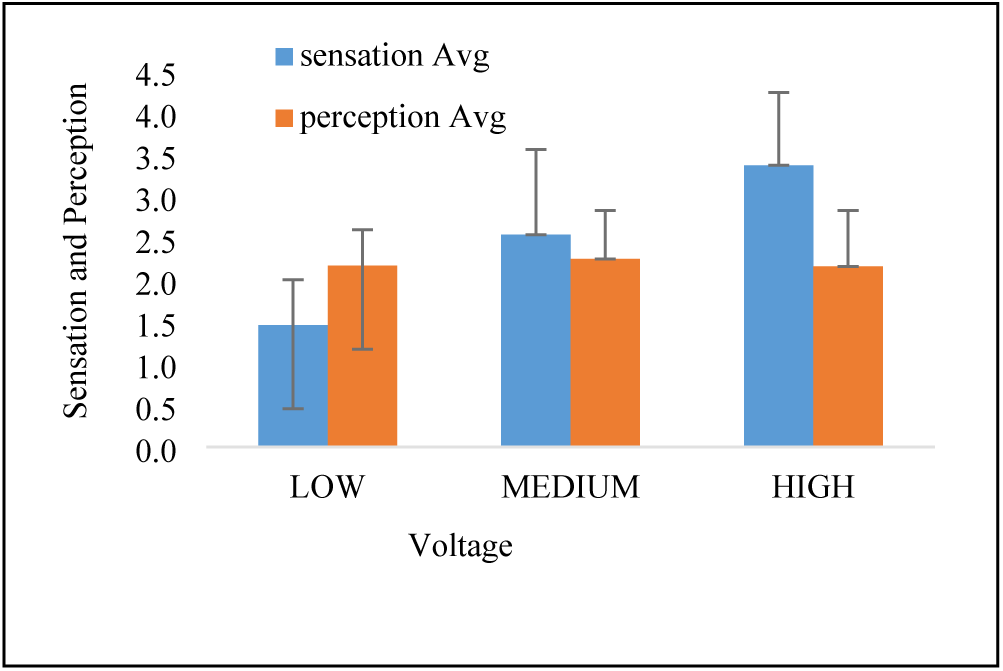
Effect of changing voltage on sensation and perception.

We see that sensation intensity increases with voltage increase, while perceived pleasantness remains stable across the different values. We calculated the average sensation and perception for a given voltage value across the 15 participants and for all the motor locations. Error bars represent one standard deviation of the sensation/perception value.

We also analyzed how sensation intensity varies depending on the location of the vibrating motor on the upper extremity (see Fig. 4). We found that sensation intensity followed similar trends across the different motor locations when changing the voltage. However, we note that the motor locations have different baseline sensation intensities, meaning that, on average, participants will report different stimuli intensity for the same voltage depending on the location of the motor on the upper extremity. We observe that the brachioradialis and the index finger are the motor locations with the highest sensation intensities, while the triceps brachii and the brachialis are the motor locations with the lowest sensation intensities. Even when the motors were placed in the same region on the upper extremity, we found some differences in sensation intensity. For example, we found that the biceps brachii long head and biceps brachii short head have a slightly higher (by 0.89 ± 0.44) sensation intensity than the posterior triceps brachii and the anterior brachialis (t(549)=13.65, p<0.0001). In the shoulder, we observe that the medial and anterior deltoids have a slightly higher (by 0.41 ± 0.03) sensation intensity than the posterior deltoid (t(413)=3.91, p<0.001).

**Fig. 4.**
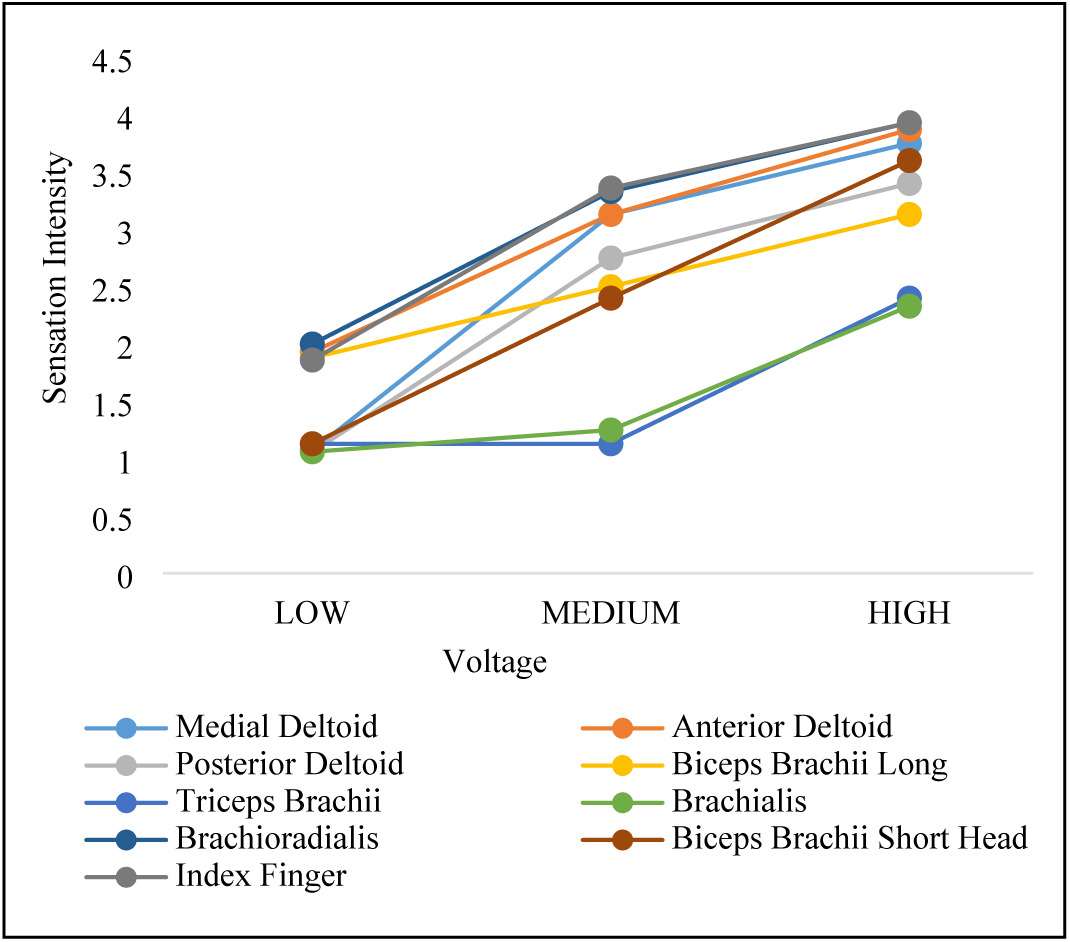
Average sensation intensity per motor location for still study.

Average sensation intensity per motor location at different values of voltage for 15 participants in the still study.

### Sensation analysis on the moving upper extremity

Our second study investigated the effects of changing voltage on sensation intensity while the participant is doing reaching movements. As described in Section IV.C, we focused on a subset of the muscles in Fig. 2, which were the medial deltoid, triceps brachii, biceps brachii short head and brachioradialis. In Fig. 5, we compare the sensation intensity in these muscles for the still and moving studies. We observe an average decrease of 5.9 ± 2.9% in sensation intensity in the moving study compared to the still study. Although the average decrease is slight, it is statistically significant (t(1726)=2.9, p<0.005). Most of the decrease occurs in the biceps brachii short head (9.6%) and the brachioradialis (6.7%). The decrease in the medial deltoid and triceps brachii are not statistically significant.

**Fig. 5.**
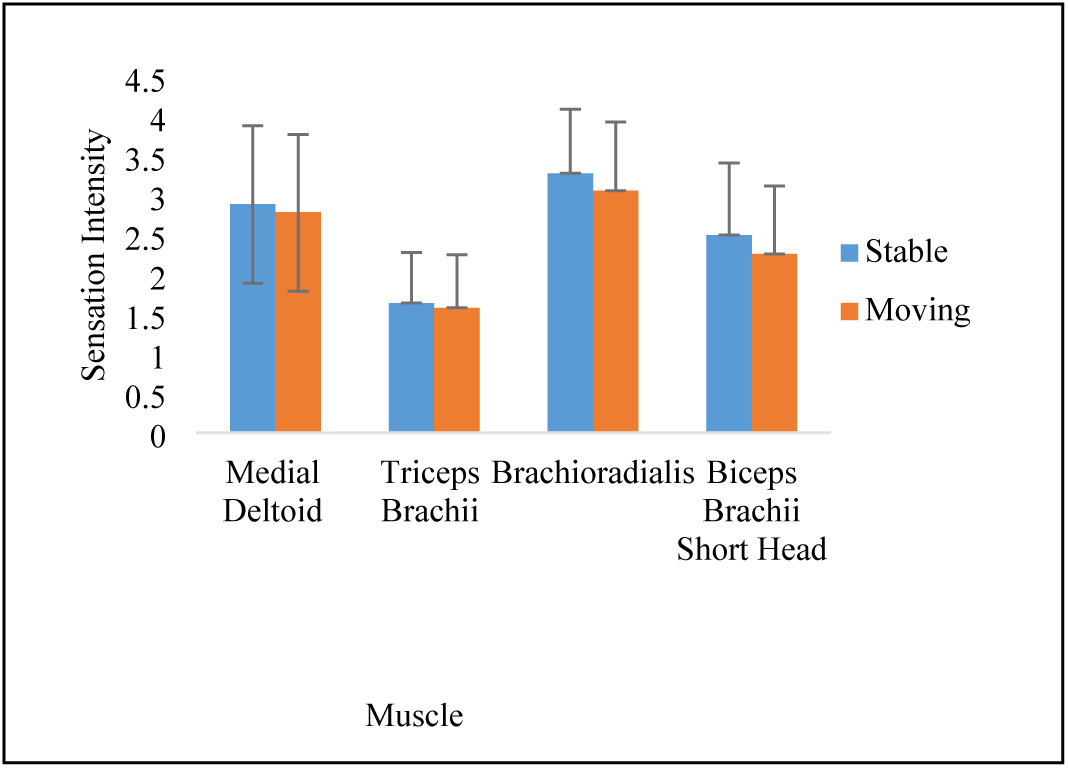
Average sensation intensity per motor location still versus moving studies.

Average sensation intensity per motor location at different values of voltage for 15 participants. We observed a decline in sensation intensity in the moving study. Error bars represent one standard deviation of the sensation intensity.

### Motor identification analysis

We analyzed the effect of increasing voltage on motor identification in both the still and the moving studies. We found that as we increase voltage, motor identification accuracy in the still study increased from 33.3 ± 16.5 % for low voltage to 80.9 ± 15.4 % for high voltage. Similarly, in the moving study, motor identification accuracy increased from 19.6 ± 22.3 % at low voltage to 89.3 ± 12.8 % at high voltage (Fig. 6). Note that we had a smaller number of stimulated muscles in the moving study compared to the still study, which makes the identification task easier. On average, participants were able to identify the right motors 72.0 ±13.8 % of the time in the still study (Fig. 7) and 72.5 ± 25.6 % in the moving study (Fig. 8).

**Fig. 6.**
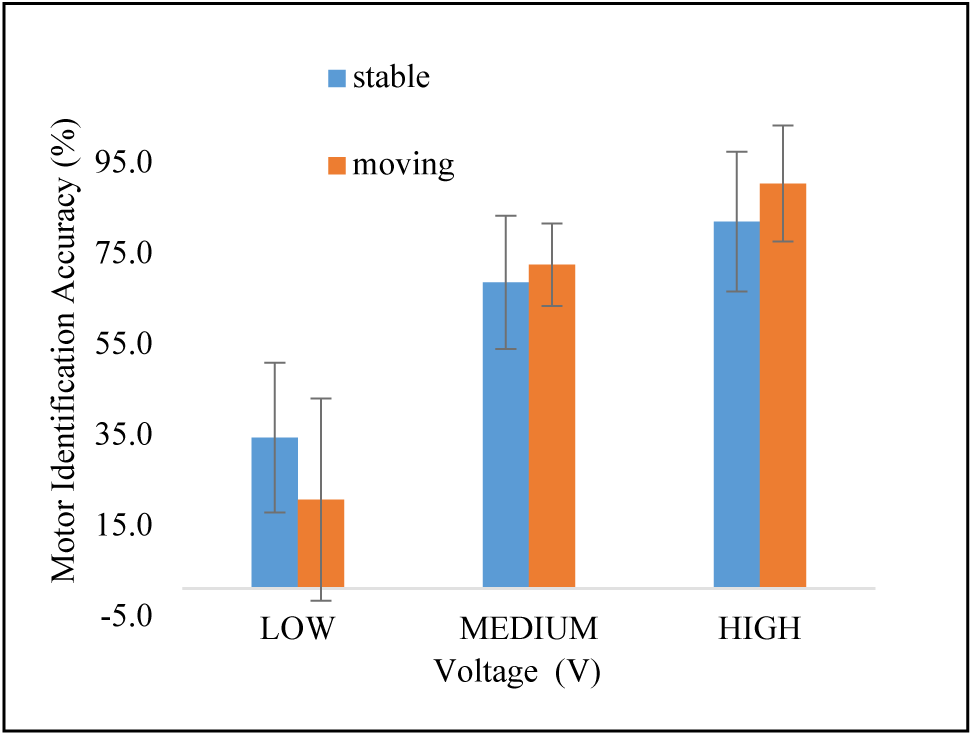
Motor identification accuracy at different voltage values in the still versus the moving studies.

**Fig. 7.**
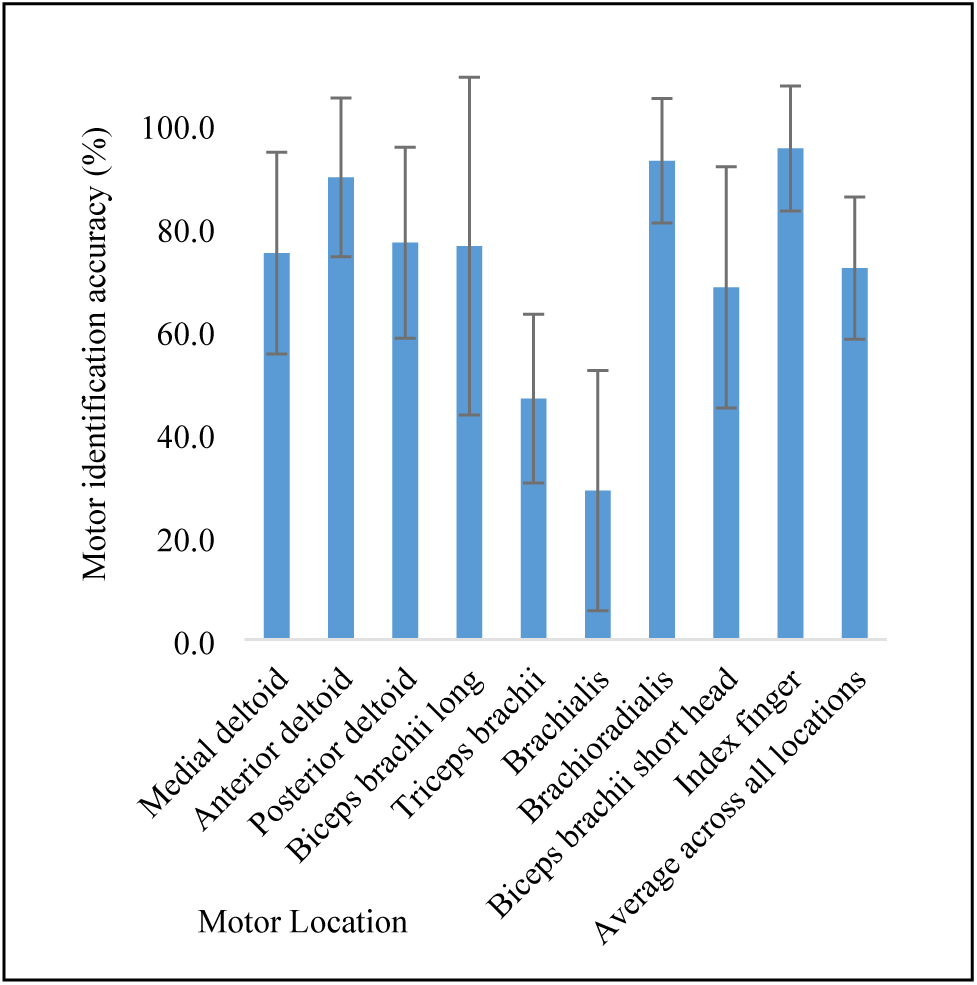
Average percentage of motor identification accuracy per motor location for 15 participants in still study.

**Fig. 8.**
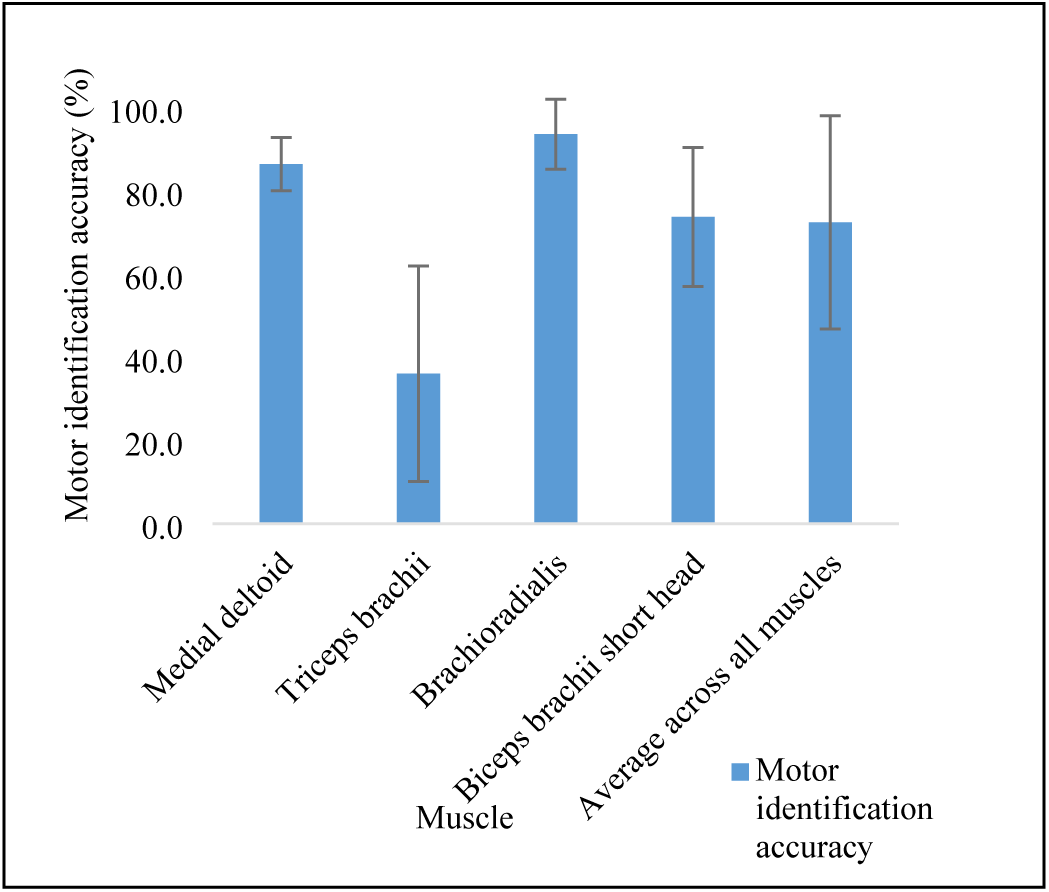
Average percentage of motor identification accuracy per muscle in the moving study.

We observe an increase in accuracy with increased voltage value.

Error bars represent one standard deviation of the identification accuracy.

We performed an analysis on motor identification per muscle in both the still and moving studies. We found that some locations such as the index finger and the brachioradialis have higher motor identification accuracy at low voltage compared to other locations such as the triceps, brachialis, and biceps brachii short head, which require higher stimuli to be clearly recognized ( Fig. 9). This was also true in the moving study, for example triceps brachii requires higher stimulation to be recognized compared to biceps brachii short head, medial deltoid, and brachioradialis (Fig. 10). Intuitively, we observe that muscles with lower sensation results (Fig. 4) have a lower motor identification accuracy.

**Fig. 9.**
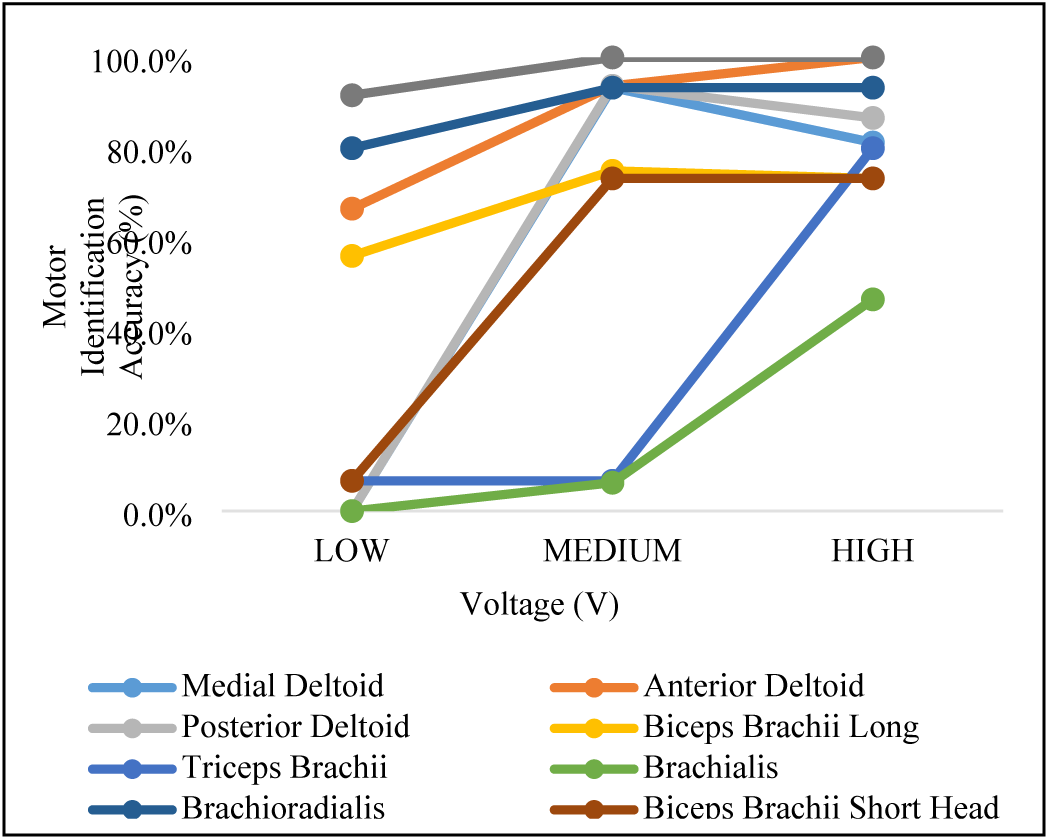
Motor identification accuracy per motor location in 15 participants in still upper extremity study over different values of voltage.

**Fig. 10.**
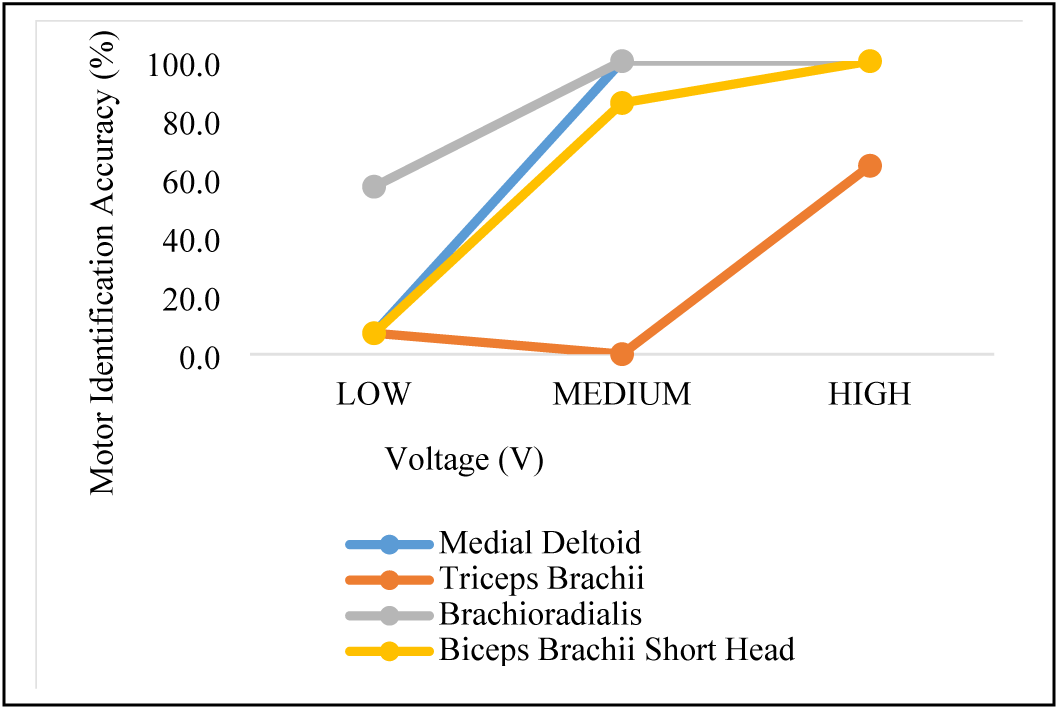
**Motor identification accuracy per muscle in the moving upper extremity study over different voltage values.**

The figure shows how different motor locations require different voltage values to reach a higher motor identification percentage.

### Multiple vibrating motors analysis

Our last study was focused on the accuracy of identifying simultaneous vibrating motors. We had three motors vibrating simultaneously with high voltage input value of 5V using four different sets of motor locations as described in sec. IV. D. As shown in Fig. 11, participants were able to identify all three vibrating motors 30.0 ± 33.0 % of the time, and two out of the three vibrating motors 41.7 ± 32.3 % of the time. Participants were only able to vaguely identify the motor location 8.3 ± 20.4 % of the time (i.e., by specifying the location as in the shoulder, upper arm or lower arm) and missed identifying all three motors 3.3 ± 8.8 % of the time. Overall, participants were able to identify the right motors 64.4 ± 19.6 % of the time. We also found that participants were able to identify an average of 2.5 ± 0.8 out of three vibrating motors.

**Fig. 11.**
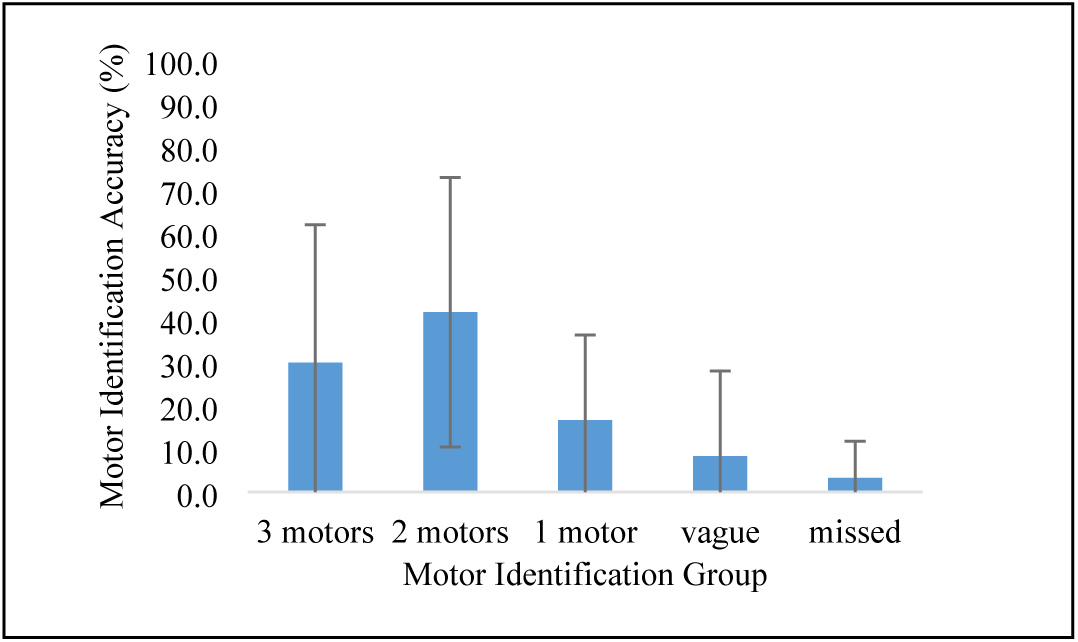
Motor identification accuracy in the three simultaneously vibrating motors study averaged over 15 participants.

We observe that most participants accurately identified only two out of the three vibrating motors. Error bars represent one standard deviation of the motor identification accuracy.

### Discussion

In the sensation study on the still upper extremity, we found that increasing vibrotactile stimuli voltage results in higher sensation intensity, with moderate to strong positive linear correlations of r=0.39. Previous work in psychophysics have noted a power law relationship between stimulus amplitude and sensation intensity. We did not observe such a relationship, perhaps because our stimuli parameters were not discretized finely enough, or their range of values was too restrictive [15][16]. In prior work, it was found that sensation intensity increases with frequencies up to 100 Hz, and then plateaus [7]. Other authors found that high amplitude and low frequency stimuli were perceived more intensely than high frequency and low amplitude stimuli[7]. We could not investigate this in this work as we have used ERM motors, where frequency and voltage are linked together.

In future work, it would also be interesting to investigate different kind of motors, such as Linear Resonant Actuators for high frequency stimulation [1], and examine the use of electrotactile feedback, where frequency and voltage can be controlled separately [13]. As for the perceived pleasantness, we did not see a significant difference (i.e., perception values stayed in the neutral range) when changing the voltage, on average.

Compared to other work in the area, Seim et al. [1] reported that subjects in both a stroke and healthy group were dissatisfied with a stimulation that they could not sense and enjoyed high voltage stimuli. We did not see a significant correlation between high voltage stimuli and perceived pleasantness on average. In our work, we found that participants on an individual level expressed different feelings with respect to the stimulation, some reporting neutral perception across the range of voltage values, others indicating a more pleasant feeling with higher voltage, and others indicating an unpleasant feeling with high voltage values. Some participants reported that the stimulation applied at certain locations on the upper extremity created a feeling of annoyance, tingling or massaging. Because we did not find a strong aggregate-level correlation between these parameters and the perceived pleasantness of the simulation, this indicates the need for customizing the parameter tuning to the individual person. One limitation of our work is that we did not vary the vibration patterns or durations. In future work, it could be interesting to investigate the effects of these parameters on perception as some subjects reported that they might have felt differently if the vibration duration was increased. Seim et al. [1] observed that participants reported a lower sensation intensity when the stimulation was applied in the lower arm (however, they did not target specific muscles in the lower arm). In contrast, we found that certain lower arm muscles such as the brachioradialis produce a high sensation intensity (Fig. 12). This finding can help in developing haptic devices and in understanding at what locations the stimulation should be applied.

**Fig. 12.**
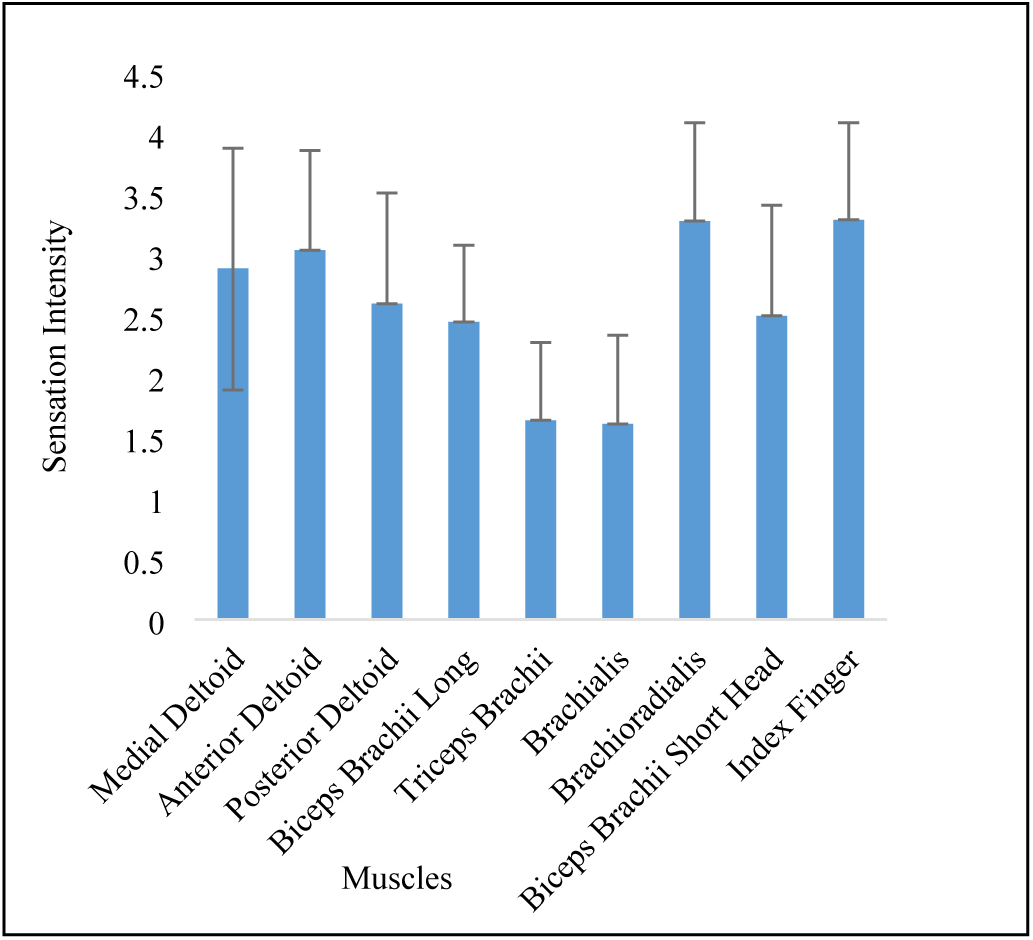
Average sensation intensity across the different motor locations averaged over the 15 participants and voltage values in the still study.

Error bars represent one standard deviation of the sensation intensity.

In the moving upper extremity study, we found that sensation intensity also increased while increasing voltage, but on average the increase is less than the measured one in the still study by 5.9 ± 2.9% (Fig. 5). Related work in the field [8] found that sensation intensity in the hand decreases as it is moving. Our work confirms that we observe the same pattern across the upper extremity. One limitation of our work is that we asked participants to perform simple reaching movements in the moving study, which will impact certain muscles in the upper extremity more than others. A topic of potential interest in future work would be to determine whether other upper extremity movements or the degree of fatigue produce significantly different sensation intensity results.

In the motor identification study, our goal was to determine whether participants were able to locate the vibrating motors in certain areas more than others, without visual feedback. We found that the brachioradialis muscle has a higher motor identification accuracy at low stimuli compared to other muscles like the triceps, brachialis, and biceps brachii short head (Fig. 9). This is consistent with our results in Fig. 4 that indicate that participants reported a higher sensation intensity to the brachioradialis than these other muscles. These differences in sensation intensity might be explained by differences in the densities of the mechanoreceptors across the upper extremity, but these have not yet been characterized [9]. Intuitively, we also found that motor identification accuracy broadly increased as we increased the voltage parameter in Fig. 10. However, for the highest voltage value we observe that the motor identification accuracy decreases for most locations. This indicates that participants start losing motor identification accuracy when the stimulus intensity of the vibrating motor increases above a threshold value. One of the limitations of our work is that participants did not have noise-cancelling headphones to prevent them from using auditory feedback to identify the motors. Another limitation might be in the hardware of the vibrating motors as we cannot guarantee that all motors give the same stimuli intensity with the same set of parameters. It is also known that ERM motors exhibit various frequencies as they accelerate and decelerate during a given activation [17]. It would also be interesting to examine other muscles and locations in the upper extremity to stimulate in addition to those examined in this work (Fig. 2).

Our final study focused on motor identification accuracy with simultaneously vibrating motors. We found that participants were able to accurately identify two out of three vibrating motors 41.7 % of the time compared to 30 % of the time for identifying all three motors. Examining the cases where only two motors were identified accurately, we observe that participants were able to more easily identify the motors that came from different regions on the upper extremity (see Fig. 13). Shah et al. [9] found that sequential vibrotactile stimuli results in better intensity discrimination than simultaneous stimuli, independent of whether the pair of motors were located within the same dermatome or across dermatomes. In our work, we found that simultaneous stimulation across the upper extremity decreases motor identification accuracy compared to single motor stimulation (Fig. 13). Bark et al. [3] found that vibrotactile stimulation can help learn simple arm movements involving 1 degree-of-freedom, but no significant effect was found in more complex movements involving 2 or 3 degrees-of-freedom. The results are consistent with our study that show that most participants can only identify 2 vibrating motors at a time, which could limit the ability to teach complex movements. Teaching complex movements would likely require more vibrating motors, which could indicate the need to investigate new vibrotactile stimulation strategies that only activate a small number of motors simultaneously, and that uses sequential vibration strategies to teach new muscle coordination patterns.

**Fig. 13.**
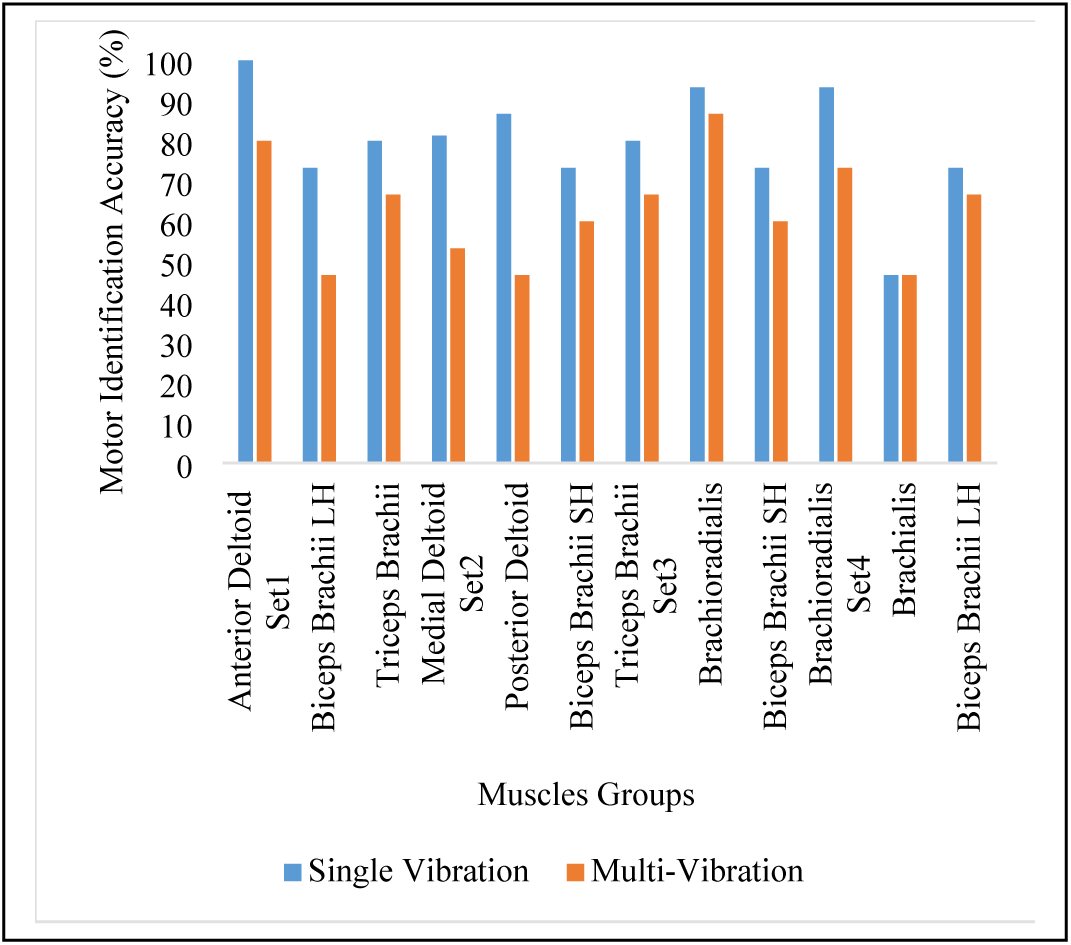
Exact motor identification percentage across the four sets of muscles.

In this figure we show the results of comparing simultaneous versus single stimuli effect on motor identification, it also shows how motor identification accuracy increased for muscles of different groups in the multi-vibrating study.

## Conclusion

We have developed a vibrotactile stimulation system designed for the upper extremity. In this work, we have conducted three studies on 15 participants to evaluate the effect of changing ERM motors stimuli voltage on sensation intensity, perceived pleasantness, and motor identification. We have found a significant correlation between voltage and sensation intensity, but no significant aggregate-level correlation between voltage and the perceived pleasantness of the simulation, which indicates the need to customize the parameters for individual subjects. This could be investigated with human-in-the-loop optimization in future work [14]. Our work identifies which motor locations in the upper extremity produce a greater sensation intensity and are easier to identify without visual feedback. We have also investigated how movements impact sensation intensity and whether subjects were able to identify simultaneously vibrating motors (without visual feedback). We expect our results to hold for other types of vibrating motors, but future experiments would need to confirm this. Our results will aid in designing haptic vibrotactile stimulation devices for the upper extremity and in developing vibrotactile stimulation patterns for applications in human-computer interaction, motor learning and rehabilitation. Our future work will focus on designing a wearable upper extremity vibrotactile stimulation device that is easy to wear and that does not obtrude the subject’s movements, which is especially important for rehabilitation studies.

